# Methylosome and SMN complexes are dispensable for snRNP assembly in Arabidopsis

**DOI:** 10.1101/2023.09.06.556483

**Authors:** Daniela Goretti, Silvio Collani, Sarah Muniz Nardeli, Markus Schmid

## Abstract

The role of RNA splicing as modulator of the molecular responses to stress is well described. In contrast, its importance in the acclimation of plants to changes in ambient temperatures started to emerge only recently. Here, we analyzed the role of temperature in spliceosome assembly, a key step often neglected in studies focusing on splicing. Taking advantage of mutants showing temperature-dependent phenotypes we conducted a comprehensive study of the role that the methylosome and SMN complexes have in plant snRNP assembly. Genetic analyses, as well as *in vivo* and *in vitro* evidence suggest a mechanism for snRNP assembly in plants that differs remarkably from vertebrate animals. The SMN complex in plants is apparently reduced to a single protein, GEMIN2, that is not essential for plant development. Similarly, the methylosome has a less crucial role in spliceosome assembly than previously thought. Our results highlight how an evolutionary conserved molecular process like RNA splicing has nevertheless evolved plant specific characteristics.

## INTRODUCTION

RNA splicing is an essential regulatory mechanism occurring in all eukaryotic organisms. The splicing reaction, during which introns are removed from the primary transcript and exons are joined, is catalyzed by large ribonucleoprotein complexes, the spliceosomes (Wahl et al., 2009). At the heart of each spliceosome are seven Sm proteins, called B/B’, D1, D2, D3, E, F, and G, that form a heptameric ring into which different U-rich small nuclear RNAs (snRNAs), U1, U2, U4, and U5, are incorporated, after which the resulting small nuclear ribonucleoprotein (snRNP) complexes are named. In addition, splicing requires a fifth snRNP that consists of a ring of seven Sm-like (LSm) proteins, LSm2-LSm8, and the U6 snRNA. An alternative heptameric LSm ring, consisting of LSm1-LSm7, is involved in cytoplasmic nonsense-mediated RNA decay (NMD) (Mayes et al., 1999; He and Parker, 2000; Golisz et al., 2013).

Importantly, in animals the assembly of the Sm and LSm rings involves different mechanisms. LSm proteins have the inherent capacity to self-assemble into ring-shaped multiprotein complexes, binding snRNAs only in a second step (Achsel et al., 1999). In contrast, Sm proteins, although originating from a duplication of LSm genes, have lost their ability to self-assemble *in vivo* and are assembled in a step-wise process (Friesen et al., 2001; Meister and Fischer, 2002; Veretnik et al., 2009)

In vertebrate animals, the assembly of the Sm ring in the cytoplasm requires two protein complexes referred to as methylosome and Survival Motor Neuron (SMN) complex (Matera and Wang, 2014). The human methylosome consists of the Protein Arginine Methyltransferase 5 (PRMT5/SIP1) and Methylosome Protein 50 (MEP50/WP45) (Antonysamy et al., 2012). These core components bind to Chloride Channel Nucleotide Sensitive 1A (PICLN/CLNS1A), which acts as molecular chaperone, to facilitate methylation of several Sm proteins (Pesiridis et al., 2009). Specifically, PRMT5 is a type II arginine methyltransferase that catalyzes the symmetric dimethylation (SDM) of RG motives in the C-terminal region of the B/B’, D1 and D3 Sm proteins (Bedford and Richard, 2005; Bedford and Clarke, 2009). This post-translational modification increases the affinity of these subunits to the SMN complex, facilitating the next step of snRNP assembly. Interestingly, in *Drosophila melanogaster*, which belongs to the order Diptera, methylation of Sm proteins by Dart5, the PRMT5 ortholog in fruit flies, is dispensable for snRNP biogenesis, indicating that although the process of RNA splicing is shared across the eukaryotic kingdoms and the Sm proteins and snRNAs are highly conserved, the requirement for Sm protein methylation has changed among taxa throughout evolution (Gonsalvez et al., 2008; Kroiss et al., 2008).

Similar to the methylosome, also the SMN complex differs between taxa. The human SMN complex includes the SMN protein (encoded by the duplicated *SMN1* and *SMN2* genes) and six GEMIN proteins (GEMIN2-GEMIN7) with different molecular functions (Gubitz et al., 2004). The SMN complex brings together the intermediate SmD1-SmD2 and SmB-SmD3 heterodimers and the SmF-SmE-SmG heterotrimer and assembles them all around a specific snRNA molecule, providing binding specificity to avoid Sm proteins from binding illicit RNAs (Pellizzoni et al., 2002). Upon interaction with import factors, the entire complex translocates into the nucleus where eventually the SMN complex disassembles, exerting other nucleus-specific functions, and the snRNP biogenesis is completed (Matera and Wang, 2014; Gruss et al., 2017). Even within the metazoan clade, the large SMN complex found in humans, and vertebrates in general, is considerably simplified in other orders. For example, in *Drosophila melanogaster*, the SMN complex consists of only GEMIN2 and SMN (Kroiss et al., 2008). The snRNP assembly complex is even further reduced in *Saccharomyces cerevisiae*, in which it consists only of the GEMIN2 ortholog BRR1P (Kroiss et al., 2008). Importantly, BRR1P is dispensable for vegetative growth of yeast cells under standard conditions as the Sm-ring can self-assemble. However, BRR1P is required in presence of specific perturbations, such as weak *sm* mutant alleles (Schwer et al., 2017). The contribution of arginine methylation to snRNP biogenesis and RNA splicing in yeast has not been studied extensively (Low and Wilkins, 2012; Hamey and Wilkins, 2023).

Interestingly, based on phylogenetic analyses, the plant SMN complex resembles the minimal complex found in *Drosophila* consisting of GEMIN2 and SMN (Kroiss et al., 2008), even though the role of the candidate plant SMN protein in snRNP assembly and splicing has not yet been tested experimentally. In contrast, the methylosome is even further reduced in plants, which lack a MEP50 ortholog, and its role in snRNP assembly is rather unclear (Kroiss et al., 2008; Mateos et al., 2023). The hypothesis that the plant methylosome consists of only PICLN and PRMT5 is supported by the finding that PRMT5 directly interacts with PICLN and Sm proteins and by the lethality of the *picln prmt5* double mutants (Huang et al., 2016; Mateos et al., 2023). However, phenotypic and RNA-seq analyses indicated that PRMT5 played only a minor role in modulating RNA splicing (Mateos et al., 2023).

An important aspect of RNA splicing common to all eukaryotic organisms is its modulation by environmental factors, such as temperature. Indeed, from humans to plants, organisms can sense changes in temperature and integrate this information at the molecular level through alternative pre-mRNA splicing (AS) (Bartok et al., 2013; Dikaya et al., 2021). Which splice acceptor and donor sites are used in the AS of a particular transcripts involves processes such as chromatin remodeling, regulation of the transcriptional machinery and kinetic, and post-translational modifications of splicing factors (Kornblihtt et al., 2013). More recently, and specifically in plants, also a core component of the spliceosome, *SME1/PORCUPINE* (*PCP*), and members of the snRNP assembly machinery, *PICLN* and *GEMIN2*, have been reported to modulate AS in response to temperature (Schlaen et al., 2015; Capovilla et al., 2018; Huertas et al., 2019; Mateos et al., 2023).

Importantly, while *SMN, GEMINs, PICLN*, and *PRMT5* loss-of-function mutants are lethal in mice, the corresponding single mutants are viable in plants (Jablonka et al., 2000; Pu et al., 2000; Monani, 2005; Tee et al., 2010). Here we took advantage of these viable mutants and the fact that their phenotypes are modulated by ambient temperature to explore the complexity of the snRNP assembly in *Arabidopsis thaliana* and its effect on plant growth and development. Overall, our analyses support the hypothesis that the SMN complex in plants consists exclusively of Gemin2 and that methylation of Sm proteins plays a minor role in snRNP assembly in plants across a wide range of ambient temperature.

## Results

### Ambient temperature modulates *GEMIN2* effects on plant growth

In Arabidopsis *GEMIN2* was shown to affect splicing in response to cold treatment and to be necessary for plant growth at 10°C (Schlaen et al., 2015). Defects in splicing were also linked to growth defects in *gemin2* mutants growing at 22°C (Schlaen et al., 2015). Aiming at characterizing the role of GEMIN2 in ambient temperature responses, we grew *gemin2-2* at temperatures ranging from 16°C and 27°C, which are considered the borders of the ambient temperature range, beyond which Arabidopsis experiences cold stress or heat shock (Guo et al., 2018; Hayes et al., 2021).

At 23°C, *gemin2* mutants exhibited pleiotropic growth defects including early flowering phenotype, smaller rosette size, reduced leaf curvature, altered leaf shape, and short petioles **(Fig. 1A,B; Supplemental Fig. S1A**,**D**,**E; Supplemental Table S1)**. All these phenotypes were strongly enhanced when the mutant was grown at low ambient temperature (16°C) **(Fig. 1A,B; Supplemental Fig. S1A**,**D**,**E; Supplemental Table S1)**. Under this condition, *gemin2-2* displayed bushy growth with crumpled leaves that were spaced irregularly compared to the spiral arrangement present in the wildtype **(Fig. 1A)**. Plants also remained very small during their entire life cycle, with a rosette diameter of less than 2 cm 70 days after sowing, at which point the plants started to bolt and set siliques (**Supplemental Table S1**). Interestingly, in *gemin2-2* part of the developmental defects were rescued when grown at elevated ambient temperature (27°C) **(Fig. 1A; Supplemental Table S1)**.

**Figure 1.**
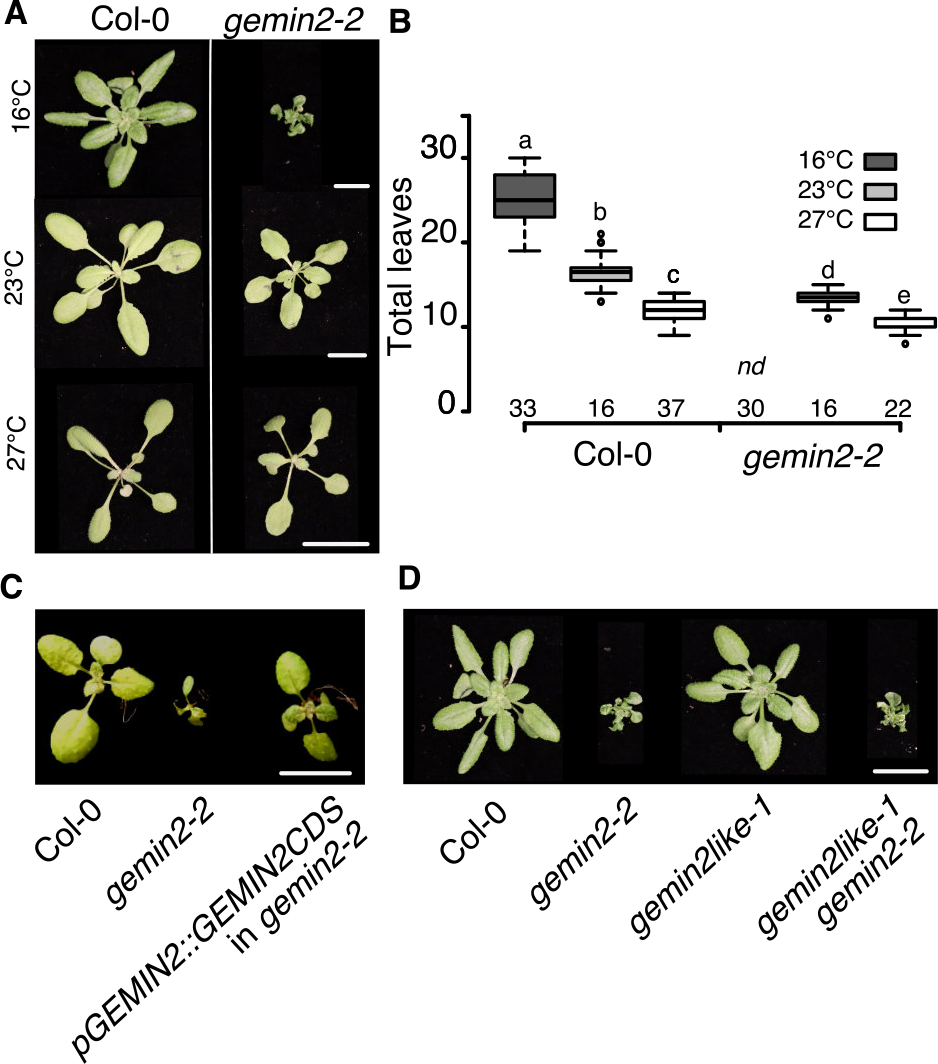
*gemin2* phenotype is modulated at ambient temperature. A, Phenotype of *gemin2-2* and Col-0 grown in LD at different temperatures. Photos are taken at 18, 25, and 33 days after sowing for plants grown at 27°C, 23°C, 16°C, respectively. B, Flowering time of Col-0 and *gemin2-2* grown in LD at three different temperatures. Numbers on the x-axis indicate the number of individuals. One-way ANOVA was used for statistical analysis. Different letters indicate categories that are statistically different (p≤0.05). C, Rescue of the *gemin2-2* mutant by a *pGEMIN2::GEMIN2CDS* transgene. Plants were grown at 16°C in LD and photos were taken three weeks after sowing. D, Comparison of the rosette phenotype of Col-0, *gemin2-2, gemin2like-1*, and *gemin2-2 gemin2like-1* double mutants grown in LD at 16°C for 33 days. Scale bars: 1 cm.w

The strong *gemin2-2* phenotype shown by plants at 16°C could largely be reverted to wildtype by expressing the *GEMIN2* CDS under the control of the *GEMIN2* promoter (*pGEMIN2::GEMIN2CDS)*, confirming the loss of this gene as the cause of the growth defects **(Fig. 1C)**. Phylogenetic analysis identified the gene At2g42510, hereafter named *GEMIN2-like*, as a likely paralog of *GEMIN2* (At1g54380) in Arabidopsis and indeed both proteins are annotated as SMN interacting proteins and harbor a SIP1 (SMN Interacting Protein 1) domain. We therefore decided to check how *GEMIN2-like* mutants performed at different ambient temperatures. Interestingly, a *gemin2-like* T-DNA insertion mutant, in which expression was reduced by 80 percent with respect to Col-0, did not show any of the phenotypes observed in *gemin2-2* and was essentially indistinguishable from wildtype (**Fig. 1D; Supplemental Fig. S1B**,**C**). To test for additive or epistatic effects between the two genes we next generated a *gemin2-2 gemin2-like* double mutant. Irrespective of the growth conditions, the double mutant always resembled *gemin2-2* single mutant, indicating no genetic interactions or functional redundancy between the two loci (**Fig. 1D; Supplemental Fig. S1D**,**E**). Taken together, these results confirmed the role of *GEMIN2* role in the modulation of plant growth in response to the temperature and extended such role to the previously unexplored range of ambient temperatures, where this gene represents an important, although not indispensable, factor modulating plant development.

### Arabidopsis misses an obvious ortholog of the human SMN protein

In vertebrates, GEMIN2 is part of a large protein complex that is reduced to just two proteins in *Drosophila*, GEMIN2 and SMN. Whether plants encode an ortholog of the human SMN is a matter of debate as phylogenetic analyses were inconclusive (Schlaen et al., 2015; Kroiss et al., 2008). We therefore decided to analyze *AtSPF30* (At2g02570), suggested to encode a possible Arabidopsis SMN protein, using genetic and molecular approaches (Schlaen et al., 2015; Kroiss et al., 2008).

First, we phenotyped two available mutants, *atspf30-1* and *atspf30-2*, at different temperatures (Kroiss et al., 2008; Romanowski et al., 2020). We observed that the *atspf30* mutants flowered earlier than wildtype, in particular at lower ambient temperature (**Fig. 2A; Supplemental Fig. S2A)**. Apart from the early flowering phenotype the *atspf30* mutants did not display any major phenotypic defects (**Fig. 2B)**. Next, we tested if AtSPF30 interacted with GEMIN2 in the heterologous yeast-2-hybrid (Y2H) system, as is the case for the human SMN and GEMIN2 proteins. However, we did not detect any growth of yeast on selective media (**Fig. 2C; Supplemental Fig. S2B)**, indicating no interaction in the heterologous host. Taken together, our results indicate that while *atspf30* mutants affect flowering time, the overall phenotype is much milder than that of *gemin2* mutants. Furthermore, the finding that the two proteins do not interact in Y2H supports the hypothesis that AtSPF30 might not function as an ortholog of the human SMN protein as previously suggested (Kroiss et al., 2008).

**Figure 2.**
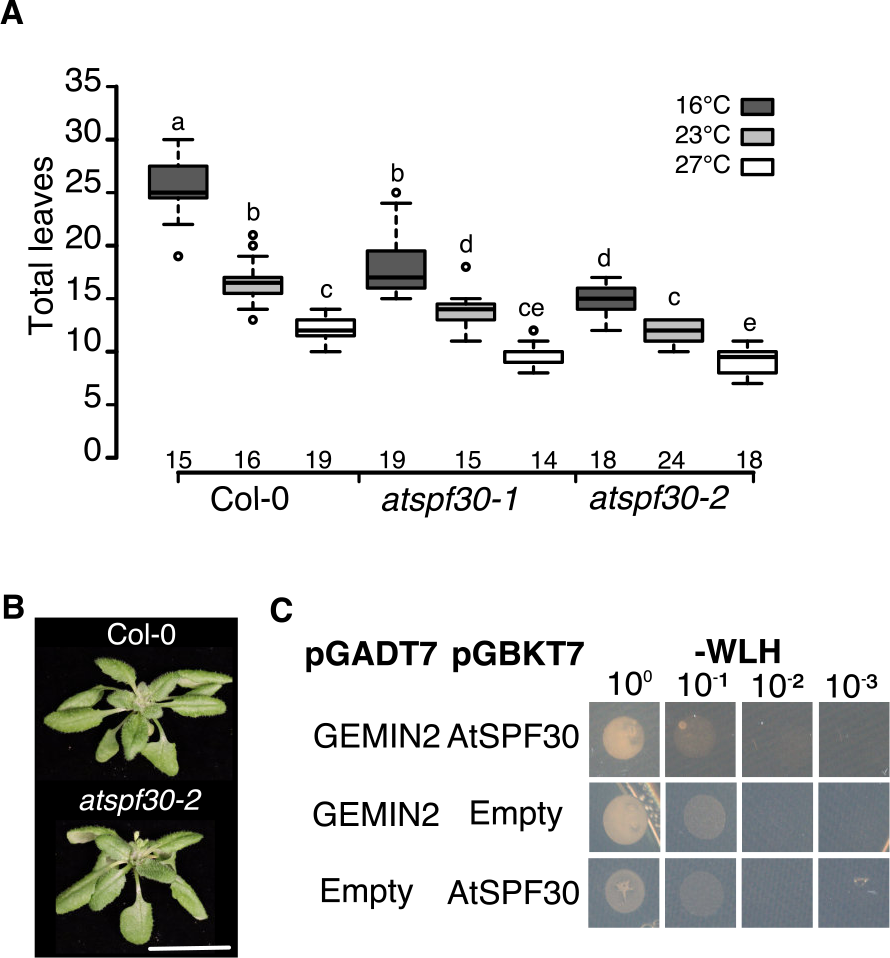
AtSPF30 is not the ortholog of the human SMN. A, Flowering time of two *atspf30* alleles and control plants grown in LD at three different temperatures. Numbers on the x-axis indicate the numbers individuals per condition. One-way ANOVA was used for statistical analysis. Different letters indicate categories that are statistically different (p≤0.05). B, Phenotype of 33-day-old *spf30-2* and Col-0 grown in LD for at 16°C. C, Interaction between GEMIN2 and SPF30. pGADT7 and pGBKT7 bring AD and BD domains, respectively. Photos were taken after four days of growth on selective drop-out medium.

Finally, to identify candidate proteins that might take over the role of SMN in the spliceosome assembly, we performed a Y2H screen using GEMIN2 as bait. Although several proteins that interacted with GEMIN2 were obtained, none of them had the expected biochemical characteristics present in the human SMN protein (**Supplemental Table S2**), such as a TUDOR domain. As neither *in silico* nor *in vitro* approaches could identify a clear SMN ortholog in Arabidopsis, we speculate that the GEMIN2 proteins in different lineages might have diversified more than anticipated and that, similar to the situation in *Saccharomyces cerevisiae*, plants might not require an SMN complex to assist in snRNP assembly.

### GEMIN2 in Arabidopsis differs from GEMIN2 in human and yeast

To test this idea, we expressed codon-optimized CDSs of human (*HsGEMIN2*) and yeast (*ScBRR1P*) *GEMIN2* in the Arabidopsis *gemin2-2* mutant. *GEMIN2* orthologs from these two species were chosen as humans form a proper multi-protein SMN complex, consisting of SMN, GEMIN2, and several other GEMIN proteins, whereas *S. cerevisiae* contains only the GEMIN2 ortholog BRR1P (Kroiss et al., 2008). We found that neither human nor yeast *GEMIN2* was able to rescue the phenotypic defects of the *gemin2-2* mutant grown at 16°C when expressed under the *AtGEMIN2* promoter (**Fig. 3A**), which is sufficient to partially rescue the mutant phenotype when expressing the *AtGEMIN2* coding region (**Fig. 1C**). In total, we analyzed more than 30 independent primary transformants (T1) per construct, all of which showed the *gemin2-2* mutant phenotype (**Fig. 3A**). To further characterize the function of GEMIN2 in plant snRNP assembly, we next tested its ability to interact with those Sm proteins that have been shown to interact with GEMIN2 in metazoans, i.e. SmE, SmF, SmG, SmD1, SmD2. Our Y2H analyses indicated that Arabidopsis GEMIN2 behaves differently from its human ortholog, interacting only with SmGa and SmGb, and to a lesser extent with SmD2a, SmD2b and SmEa (**Fig. 3B; Supplemental Fig. S3**).

**Figure 3.**
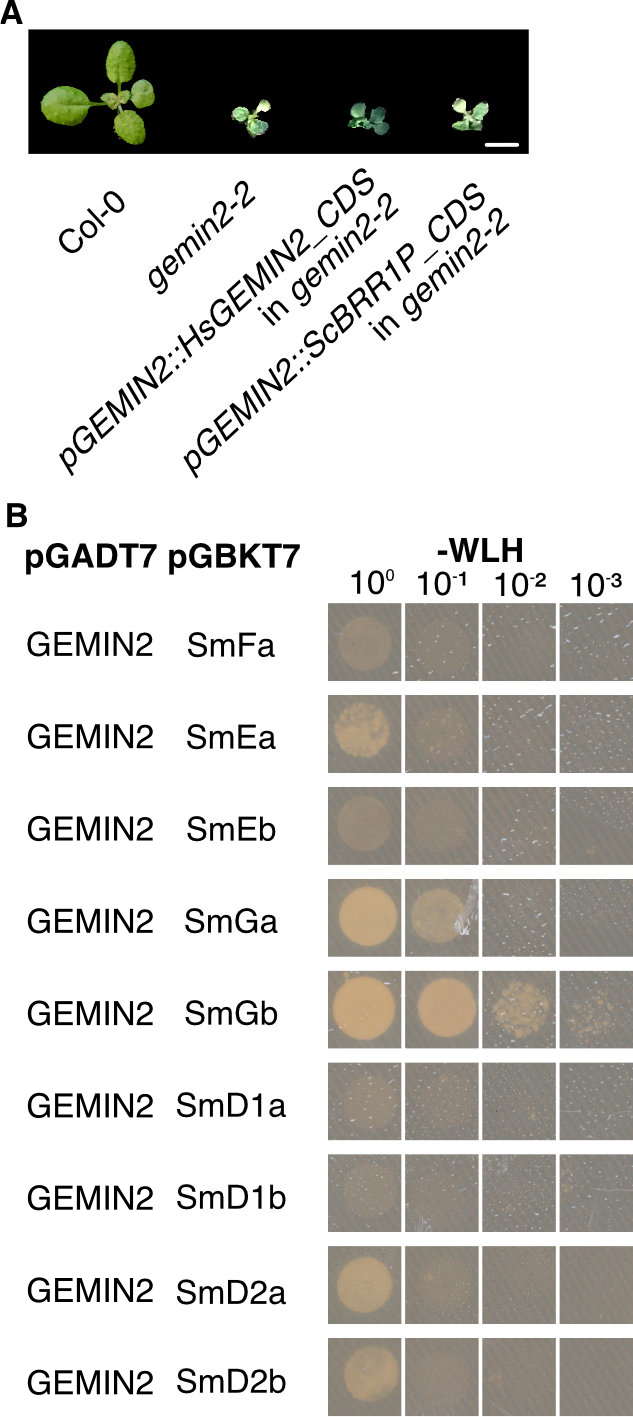
Plant GEMIN2 differs from the human ortholog. A, Phenotype of Col-0, *gemin2-2*, and of *gemin2-2* completed with human (Hs) and yeast (Sc) GEMIN2 orthologs expressed form the Arabidopsis *GEMIN2* promoter 16°C. Scale bar: 0,5 cm. B, Yeast-2-hybrid testing interaction between GEMIN2 and Sm proteins. pGADT7 and pGBKT7 vectors contribute AD and BD domains, respectively. Photos were taken after four days of growth on selective drop-out medium.

In summary, our results suggest that the plant GEMIN2 protein might differ in its function from that of its human and yeast counterparts. How exactly the GEMIN2 proteins in the different taxa acquired partially different modes of action remains to be established.

### The methylosome is not essential for plant growth and development

Having established that the metazoan SMN complex is likely reduced to just GEMIN2 in plants, we next focused our attention on the methylosome, which controls the methylation of certain Sm proteins and thus modulates snRNP assembly. In Arabidopsis, two proteins, PRMT5 and PICLN, form the methylosome and have been implicated in plant response to abiotic stresses, such as salt and cold (Hu et al., 2017; Mateos et al., 2023).

Given our results on *GEMIN2*, we decided to assess the role that PRMT5 and PICLN have in acclimation to low and high ambient temperature. We found that the *picln* and *prmt5* mutants have opposing flowering time phenotypes. *prmt5-1* (and *prmt5-2*) flowered consistently later than Col-0 at all temperatures tested, whereas *picln-2* was early flowering at 16°C but flowered approximately as wildtype at 23° and 27°C **(Fig. 4A,B; Supplemental Fig. S4A**,**D**). *picln* mutants showed growth defects especially at 16°C and have an overall smaller size than wildtype at all the tested temperatures (**Supplemental Fig. S4B**,**C; Supplemental Table S1**). Furthermore, expression of both *PICLN* and *PRMT5* is moderately induced at 16°C when compared to 23°C and 27°C, in contrast to GEMIN2, which is stably expressed across the examined temperature range **(Supplemental Fig. S4E)**.

**Figure 4.**
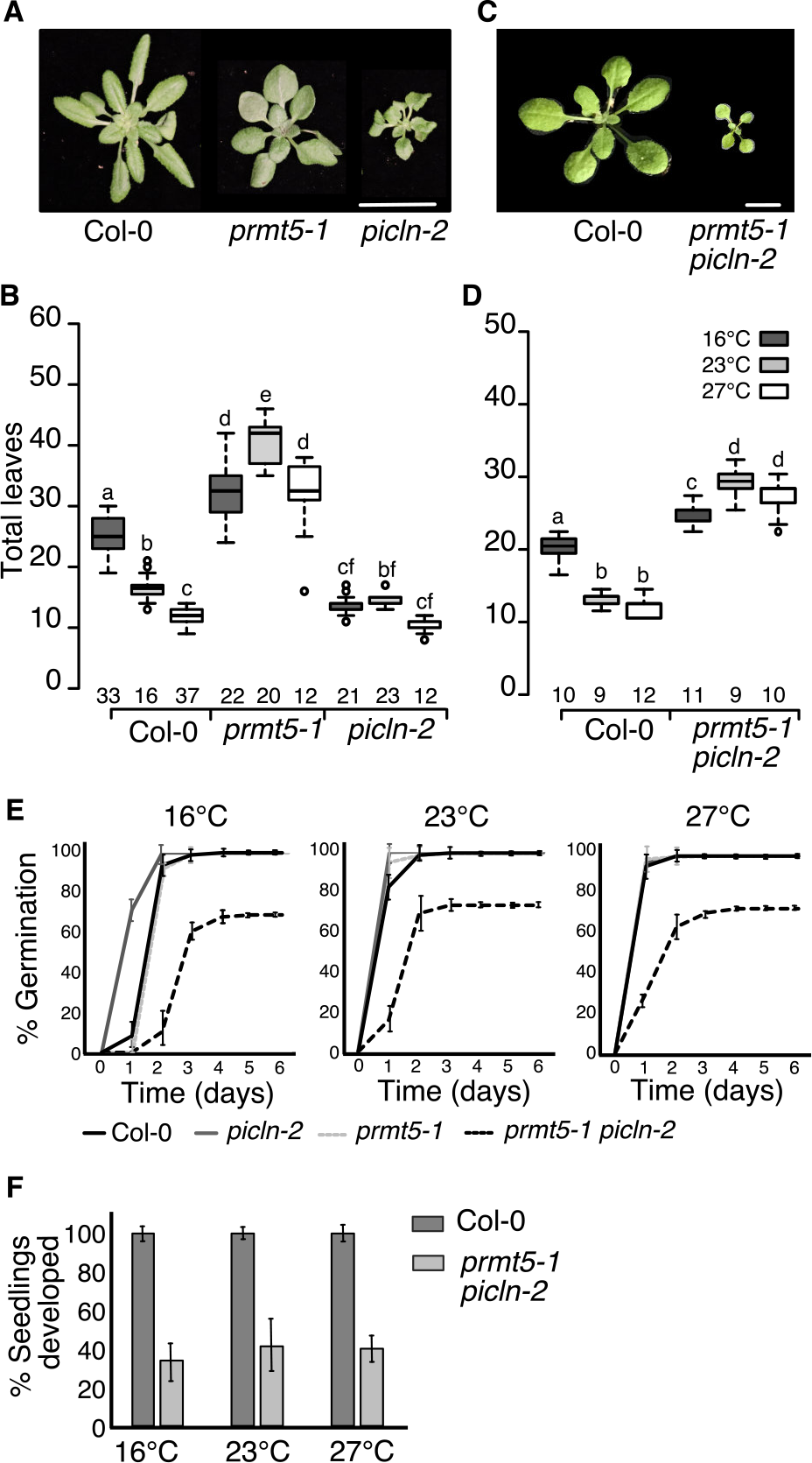
The methylosome is dispensable for plants growing at ambient temperature. A, Phenotypes of Col-0, *prmt5-1*, and *picln-2* grown in LD at 16°C for 33 days. Scale bar: 1 cm. B, Flowering time of the three genotypes in A) at three different temperatures. Numbers on the x-axis indicate the number of individuals per genotype and condition. One-way ANOVA was used for statistical analysis. Different letters indicate categories that are statistically different (p≤0.05). C, Phenotype of Col-0 and the *prmt5-1 picln-2* double mutant grown at 23°C for 20 days-after-sowing. Scale bar: 0,5 cm. D, Flowering time of the two genotypes in C) at three different temperatures. Numbers on the x-axis indicate the number of individuals. One-way ANOVA was used for statistical analysis. Different letters indicate categories that are statistically different (p≤0.05). E, Germination rate of Col-0, *picln-2, prmt5-1*, and *prmt5-1 picln-2* seeds on MS plates at three different temperatures. f, Percentage of germinated Col-0 and *prmt5-1 picln-2* seeds developing into young plants at 16°C, 23°C, and 27°C.

The opposing phenotypes of *picln* and *prmt5* mutants under the tested conditions prompted us to test the hypothesis that other proteins belonging to the PRMT family could be part of the methylosome in plants. In particular we focused on PRMT7/PRMT16 as the human homolog has previously been shown to interact with SmD3 and SmB (Gonsalvez et al., 2007; Zurita-Lopez et al., 2012). However, we were unable to detect interactions between AtPRMT7 and PICLN or any of the Sm proteins tested by Y2H (**Supplemental Fig. S5A**). Furthermore, under the tested conditions the *prmt7-1* mutant has no discernible phenotype (**Supplemental Fig. S5B**). This suggests that PRMT5 is the main arginine methylase able to interact with the splicing machinery in plants. This is in agreement with a previous report by Mateos et al. (2023) that showed that PRMT5 interacts directly with PICLN and demonstrated that the double mutant was embryo lethal, indicating genetic interaction between the two genes as already suggested by Huang et al. (2016). However, our genetic analysis doesn’t point to an epistatic interaction between PRMT5 and PICLN, suggesting that double mutants should not necessarily be lethal.

Indeed, we succeeded in establishing the *prmt5-1 picln-2* double mutant provided that the parental plants used for the cross (F0) and subsequent generations (F1; F2) were grown at 27°C, a growth conditions under which the *picln-2* phenotype is largely reverted to normal. Interestingly, the progeny of *prmt5-1 picln-2* double mutants propagated at 27°C, although generally smaller than the single mutants and Col-0 controls, were viable and set seeds even when grown at lower temperatures (**Fig. 4C,D; Supplemental Fig. S4F**). In addition, the germination rate of the double mutant was reduced by approx. 30% when compared to the single mutants and Col-0 (**Fig. 4E**). Development of 60% of the *prmt5-1 picln-2* seedlings arrested shortly after germination and plants failed to form proper leaves (**Fig. 4F**), indicating that the previously reported lethality of the *prmt5-1 picln-2* double mutant is a medium penetrance phenotype.

Taken together, the phenotype of the mutants and genetic analyses, in particular the fact that the *prmt5-1 picln-2* double mutant is viable, suggest that the methylosome might play a less prominent role in modulating plant growth and development across the ambient temperature range than previously thought.

### SDM of plant Sm proteins is dispensable at ambient temperature

It is well established that methylation of SmD1, SmD3, and SmB by the methylosome supports the assembly of snRNPs by the SMN complex in humans (Matera and Wang, 2014). However, the finding that the *prmt5-1 picln-2* double mutant is viable provides indirect evidence that the PICLN-PRMT5 methylosome is dispensable for plant development. We therefore decided to investigate the importance of symmetric demethylation (SDM) of Sm proteins for snRNP assembly and function more directly. To gain insight into the importance of SDM of SmD1b for snRNP assembly and function in plants, we cloned a truncated version of the SmD1b ORF that misses the last 16 amino acids (hereafter named SmD1bΔC) and a construct in which we mutated the last eight arginine residues in the C-terminal half of SmD1b that are potential targets for symmetrically dimethylation by PRMT5 (Brahms et al., 2001) into lysine (hereafter named SmD1b^Lys^) (**Supplemental Fig. S6A**). It should be noted that in human the equivalent of the single RG motive at position 61, which is maintained in our constructs, has been shown to not be involved in the interaction between methyltransferases and the SNRPD1 protein (Friesen et al., 2001). Therefore, we excluded this single RG from our targeted mutagenesis approaches. Y2H analyses confirmed that PRMT5 interacts with wild-type SmD1b, as already shown by Mateos et al. (2023). This interaction was abolished in SmD1bΔC and SmD1b^Lys^, indicating that the arginine residues in the C-terminal region of SmD1b were essential for the interaction with PRMT5 and also suggesting that the remaining single RG in position 61 was not sufficient to facilitated interaction between SmD1b and PRMT5 (**Fig. 5A; Supplemental Fig. S6B**). In contrast, deletion or mutation of the C-terminal region had only a neglectable effect on the interaction with SmD2a and SmD2b (sharing the same amino acid sequence), which interact with SmD1b to form one of the heterodimers from which snRNPs are assembled (Zhang et al., 2011) (**Fig. 5A; Supplemental Fig. S6B**). Taken together, these results suggest that SMD (of SmD1b) might not be required for snRNP assembly in plants.

**Figure 5.**
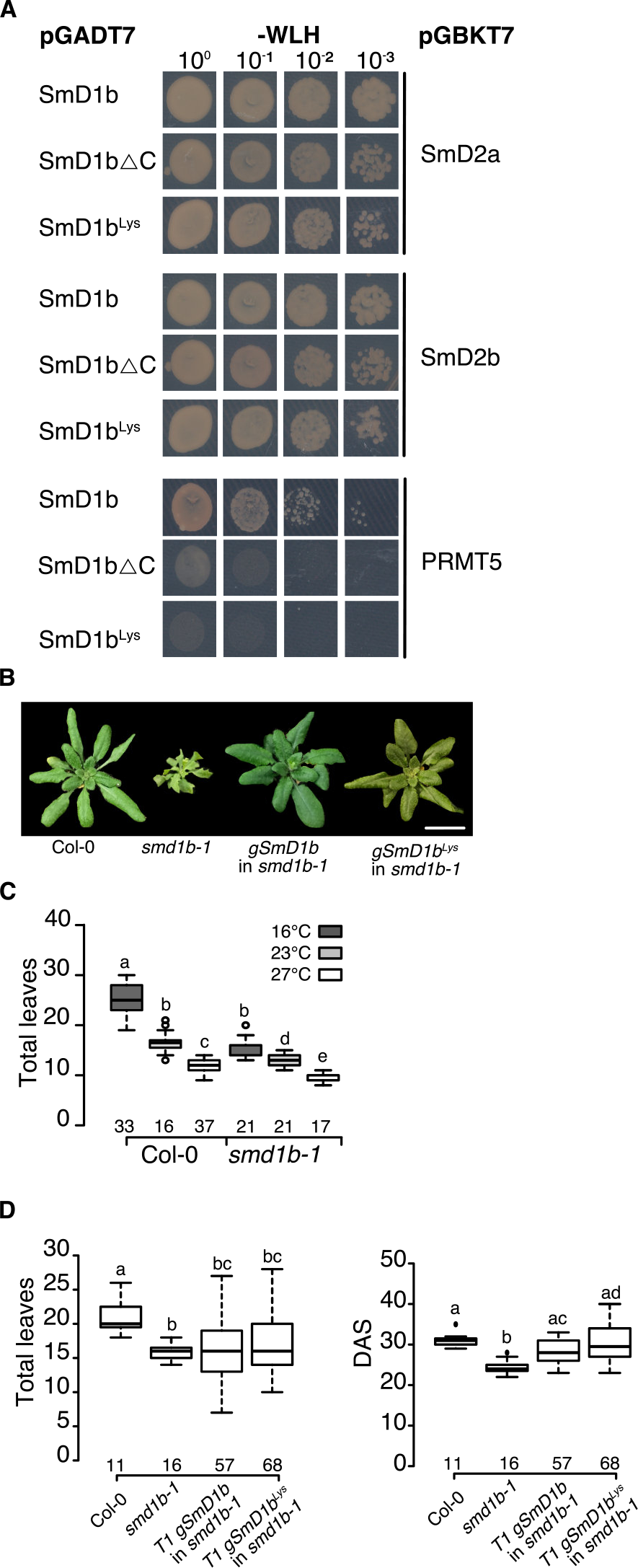
SDM of SmD1a is dispensable at ambient temperatures. A, Interaction between unmodified SmD1b, SmD1b lacking the C-terminal 16 amino acids (ΔC) and SmD1b in which 8 arginine residues in the C-terminal half have been replaced by lysine (Lys) with SmD2 and PRMT5. pGADT7 and pGBKT7 contribute the AD and BD domains, respectively. Photos were taken after four days of growth on selective medium. B, Phenotype of Col-0, *smd1b-1*, and T1 *smd1b-1* plants complemented with either wildtype (gSmD1b) or a protein version in which the last 8 arginine residues have been replaced by lysine (gSmD1b^Lys^) grown at 16°C for 30 days. Scale bar: 1 cm. C, Flowering time of Col-0 and *smd1-b* mutant at three different temperatures. D, Flowering time of Col-0, *smd1b-1*, and two T1 lines carrying either the wildtype or mutagenized version of *gSmD1b* grown at 16°C. Numbers on the x-axis indicate the number of individuals per genotype and condition. One-way ANOVA was used for statistical analysis. Different letters indicate categories that are statistically different (p≤0.05).

To test this hypothesis, we took advantage of the *smd1b-1* mutant, which displays developmental defects under normal growth conditions (Elvira-Matelot et al., 2016). Similar to the other mutants discussed, we found that the pleiotropic phenotype of *smd1b-1* was enhanced at lower ambient temperatures **(Fig. 5B; Supplemental Fig. S6C)**. At 16°C, *smd1b-1* plants were early flowering and showed altered leaf morphology and smaller rosette diameter compared to wildtype, similar to the *picln-2* and *gemin-2* mutants **(Fig. 5B,C; Supplemental Fig. S6C**,**D; Supplemental Table S1)**.

Next, we expressed the wild-type SmD1b and the mutagenized SmD1b^Lys^ ORF under the control of the 35S promoter or a 1578bp *SmD1b* promoter fragment in the *smd1b-1* background. Phenotypes were assessed in T1 plants grown at 16°C, a condition under which the mutant has the strongest phenotype. While the expression of the two proteins under the endogenous *SmD1b* promoter was not able to complement the *smd1b-1* phenotype (> 40 T1 plants), the use of the constitutive 35S promoter resulted in full complementation, indicating that the rescue construct using the *SmD1b* promoter was missing important *cis* regulatory elements (**Supplemental Fig. S6E**,**F)**. To overcome this problem, we cloned a genomic *SmD1b* wildtype rescue construct (*gSmD1b*), consisting of the 1578bp promoter, all exons, introns and a 2003bp terminator. Next, we mutated the last eight arginine residues in the C-terminal half of SmD1b in the *gSmD1b* construct, resulting in gSmD1b^Lys^. Both the wild-type and the gSmD1b^Lys^ genomic sequences complemented the smaller size and the early flowering time of the *smd1b-1* mutant in T1 to a large extent **(Fig. 5D; Supplemental Fig. 7A**,**B)**. However, around 32% of the plants transformed with the *gSmD1b*^*Lys*^ construct retained some of the developmental defects present in *smd1b-1*, such as flattened leaves and irregular leaf margins (**Supplemental Fig. 7A**,**B)**. The same phenotype was detectable in only 9% of the T1 plants expressing wildtype *gSmD1B* (**Supplemental Fig. S7A**,**B)**. The finding that a version of SmD1b that can no longer be methylated by PRMT5 mostly retains its function indicates that Sm proteins methylation is not essential in Arabidopsis, supporting the notion that the molecular mechanism controlling snRNP assembly differ in plants and humans.

## DISCUSSION

RNA splicing is of fundamental importance in plant-environment interaction, being one of the first steps enabling plants to adjust their transcriptome and proteome in response to external stimuli. In metazoans the stepwise assembly of snRNPs, which form the heart of the spliceosome, is facilitated by different protein complexes, the methylosome and the SMN complex. The core components of these complexes are evolutionary conserved, giving rise to the idea that snRNP assembly follows the same basic rules in other eukaryotic organisms, including plants (Matera and Wang, 2014; Dikaya et al., 2021). However, more recently interesting differences between snRNP assembly in plants and metazoans have emerged. It has been reported that in plants components of the methylosome and SMN complex such as PICLN, PRMT5 and GEMIN2 directly participate in the modulation of plant growth under certain environmental conditions (Pei et al., 2007; Zhang et al., 2011; Fu et al., 2013; Schlaen et al., 2015; Hu et al., 2017; Mateos et al., 2023). In this study, we focused on the role these genes play in modulating Arabidopsis development across the ambient temperature range from 16°C to 27°C, conditions in which all the tested mutants are viable.

Interestingly, we found that the phenotype of the *gemin2* mutant, which was previous shown to be seedling lethal at 4°C (Schlaen et al., 2015), is actually strongly temperature-depended. At 16°C, the mutant exhibits strong developmental defects but is viable, and the phenotype is progressively rescued with increasing temperatures. Furthermore, we were unable to detect interaction between GEMIN2 and AtSPF30, which has been suggested to function as the SMN protein in plants, and Y2H screens failed to isolate any candidate genes that resemble components of the metazoan SMN complex. While these are strictly speaking negative results, they nevertheless argue that in plants the function of the SMN complex is reduced to just GEMIN2.

While it might seem surprising that core molecular machineries such as the snRNP assembly might differ so much between taxa, the situation in plants is not unheard of as also the yeast SMN complex has been reduced to a single gene, BRR1P (Kroiss et al., 2008). However, this is not a simple case of convergent evolution as BRR1P, similar to HsGEMIN2, is unable to rescue the Arabidopsis *gemin2* mutant, indicating that the proteins have diverged throughout evolution and have acquired specific functions. It is thus not surprising that in Y2H GEMIN2 from Arabidopsis interacted only with a subset of the Sm proteins (SmGa, SmGb, SmEa) predicted based on its human counterpart. However, these protein-protein interactions directly link the plant GEMIN2 to Sm proteins, confirming its role in snRNP assembly. Taken together, our results strongly suggest that in plants the SMN complex is reduced to just GEMIN2 and that its function is dispensable under the conditions tested. Furthermore, our findings argue that even though the key players are evolutionary conserved, the mechanisms controlling snRNP biogenesis in plants differ from that in other taxa.

Further evidence for this notion comes from our analyses of the plant homologs of the methylosome components PICLN and PRMT5. In metazoan, SDM of certain Sm proteins is essential for snRNP assembly. We therefore conducted a detailed phenotypic analysis of the *prmt5, picln*, and *smd1b* mutants at different temperatures. *SmD1b* was selected as it encodes a predicted methylosome target and a knock-out mutant of the gene was available (Elvira-Matelot et al., 2016). As in the case of *gemin2-2*, the pleiotropic phenotypes of *picln* and *smd1b* at 23°C became even more pronounced at the low ambient temperature but were largely rescued at 27°C. In contrast, the phenotype of *prmt5-1* was similar across the entire temperature range investigated.

The findings that *picln* and *prmt5* mutants had opposite effects on plant development and that *prmt5* was not affected by temperature, despite its predicted role in SDM of Sm proteins and thus presumably snRNP assembly, suggested two possible scenarios. First, Arabidopsis might have a methylase that acts redundantly with PRMT5, or alternatively, SDM of Sm proteins might play a negligible role in snRNP assembly in plants. The first alternative seems unlikely as PRMT7 did not interact with PICLN and any of the Sm proteins tested in Y2H, indicating that PRMT5 might be the main methylase capable of methylating Sm proteins in Arabidopsis. We therefore decided to test the importance of SDM of Sm proteins for snRNP assembly and function using a combination of genetic, mutant complementation, and protein-protein interaction analyses. Our findings indicate that PRMT5 function is dispensable under the conditions tested. However, it has previously been reported that PRMT5 and PICLN interact directly and that the double mutant is lethal, which argues against our findings and for a key role of Sm methylation in snRNP assembly (Huang et al., 2016; Mateos et al., 2023).

Regarding their interaction, it is worth mentioning that PRMT5 and PICLN participate in the DNA damage response in human cells independently of MEP50 (Owens et al., 2020). As MEP50 is not conserved in plants, it seems possible that a PRMT5 – PICLN complex might have a similar function in plants, although this theory needs further investigation. Furthermore, RNA-seq data from Mateos *et al*. (2023) indicated both overlapping and separate molecular functions for PICLN and PRMT5 and showed that the splicing profile in *prmt5* plants differed substantially from that of *picln* and *gemin2* mutants, which were more similar to each other. Although *picln* and *prmt5* showed such different molecular profile, Mateos et al. (2023) found that both mutants flowered late, which contrasts with our finding that under our growth conditions *picln* is an early flowering mutant. A possible explanation for these differences could come from the growth conditions, as Mateos et al. (2023) grew plants under a 12h light /12h night regime whereas we used long-day photoperiod (16h light / 8h night).

Given these conflicting results we decided to test the hypothesis that the PRMT5 – PICLN methylosome might not be essential for snRNP assembly and splicing more directly by using a genetic approach. By growing plants at 27°C, under which the *picln-2* mutant growth defects are largely suppressed, we succeeded in establishing the *picln-2 prmt5-1* double mutant, previously reported to be lethal. Interestingly, once established, the *picln-2 prmt5-1* double was viable at 16°C and 23°C, even though the phenotype was more severe than that of the single mutants. However, germination and seedling development was strongly affected in the double mutants, indicating that even though these genes might not essential, they nevertheless are important for proper plant development. Taken together, our various analyses point towards a minor role for the PICLN – PRMT5 complex in plant snRNP assembly compared to metazoans, where the importance of each protein is highlighted by the lethality of the single mutants. Our analyses further suggest that the regulation of snRNP assembly and RNA splicing in general in plants differs substantially from that of metazoans.

Finally, it should be noted that the phenotypes of many splicing related mutants have recently been shown to be sensitive to temperature. Most of these mutans are cold sensitive but more recently a warm sensitive *lsm7* knock-down mutant has been reported, indicating the essential role of RNA splicing in acclimation across the entire temperature range (Nardeli et al., 2023). How and why these mutants, which are supposedly participating in the same molecular process, RNA splicing, have such different phenotypes is not understood. Clearly, more work is needed to disentangle the role of RNA splicing in temperature acclimation and whether it constitutes a practical target for enhancing plant temperature tolerance.

## MATERIALS AND METHODS

### Plant material, growth conditions, and ambient temperature phenotyping

All genotypes used in this research were in Columbia background and mutants were obtained from Nottingham *Arabidopsis* Stock Center (NASC) or provided by colleagues: *gemin2-1* (SALK_142993), *gemin2*-2 (SAIL_567_D05), *prmt5-1* (SALK_065814), *prmt5-2* (SALK_095085), *picln-1* (GABI675_D10), *picln-2* (SALK_050231), *gemin2like-1* (SAIL_691_C12), *atspf30-1* (SALK_052016), *atspf30-2* (SALK_081292), *prmt7-1* (SALK_039529), *smd1b-1* (*sgs14* mutant, from Elvira-Matelot et al., 2016). All mutants were genotyped by PCR using the primers listed in **Supplemental Table S3**. Seeds were stratified at 4°C in the dark for 72 h before being sown on soil. Plants were grown under long-day conditions (LD), 16h light/8h dark, in Percival chambers equipped with full range white LED illumination at an intensity of 120-150μmol*m^-2^s^-1^, with controlled humidity (RH 70%) at different constant ambient temperatures (16°C, 23°C and 27°C). Flowering time was measured as the number of days until the floral buds appeared; rosette diameter was measured after bolting once the stem was 1 cm long. Germination and survival rates of Col-0, *prmt5-1, picln-2* and the *prmt5-1 picln-2* double mutant were estimated by sowing seeds on half-strength MS (without sugar) agar plates.

### DNA vectors and plant transformation

With the exclusion of the sequences listed in **Supplemental Table S4**, that were synthesized by Eurofins, the sequences were amplified by PCR from cDNA or genomic DNA and cloned into GreenGate entry vectors by using the GreenGate system (Lampropoulos et al., 2013). Final constructs were assembled in pGREENII-based GreenGate plant binary destination vector pGGZ003 and transformed into *Agrobacterium tumefaciens* (strain GV3101 pMP90 pSoup) by electroporation (Gene Pulser Xcell system). Arabidopsis plants were transformed by the floral dip method. BASTA selection (0.1%, v/v) was used for screening the transgenic lines on soil. Lists of the PCR primers used for cloning and the vectors generated in this study can be found in **Supplemental Table S3 and S5**.

### Y2H assay and library screening

For testing protein-protein interactions in a one-to-one Y2H assay, the coding sequences of the tested genes were cloned into the yeast vectors pGADT7 or pGBKT7 readapted to the GreenGate cloning system (Zacharaki et al., 2022). Oligos used in the cloning are listed in **Supplemental Table S3**. Pairs of vectors, including negative controls, were used to co-transform yeast strain AH109 and colonies carrying both vectors were selected on SD medium without tryptophan (-W) and leucine (-L) (TaKaRa 630317) at 28°C. After 6 days, protein-protein interactions were tested by growing serial dilutions on SD drop-out medium lacking tryptophan (-W), leucine (-L), and histidine (-H) (TaKaRa 630319).

For the Y2H Library Screening, a cDNA expression library was generated from mRNA pooled from the aerial part of Col-0 seedlings grown at 16°C, harvested 11 and 13 days after sowing, 3, 7, and 10 hours after daylight started. The mRNA was used create a Gateway compatible cDNA entry library by using of the CloneMiner cDNA library Construction Kit (Invitrogen) according to the manufacturer’s instructions. The final cDNA library in the pDEST22 vector had a titer of 4.9×10^6^ cfu·ml^−1^. The *GEMIN2* CDS was cloned into the bait pDEST32 vector. The cDNA library screening was performed as described by de Folter and Immink (2011).

### Reverse Transcription quantitative PCR (RT-qPCR)

Total RNA was extracted using the Qiagen Plant RNeasy kit and treated with RNase-free DNaseI (Thermo Scientific) to remove DNA contamination. Complementary DNA (cDNA) was synthesized using the RevertAid First Strand cDNA Synthesis kit (Thermo Fisher) in accordance with the manufacturer’s instructions. RT-PCR was carried out using a CFX96 Real-time System (Biorad). and SYBR Green Master Mix (Bioline). The relative expressions were calculated using the 2^(−ΔΔCT)^ method. For each sample, three biological and three technical replicates were used. Primers used in RT-qPCR are listed in **Supplemental Table S3**.

## AUTHOR CONTRIBUTIONS

DG and MS conceived the project and designed the experiments; DG, SMN and SC performed the experiments and analyzed the data; DG and MS wrote the manuscript with inputs from all the authors.

## ACKNOWLEDGMENTS

This work was supported by a grant from the Knut och Alice Wallenbergs Stiftelse (KAW 2018.0202) to MS. The authors acknowledge Dr. Julieta L. Mateos for sharing seeds of *gemin2-1* and *picln-1* mutants and Prof. Hervé Vaucheret for sharing seeds of *smd1b-1* mutant.

