## Supplemental files for "Methylosome and SMN complexes are dispensable for snRNP assembly in Arabidopsis"

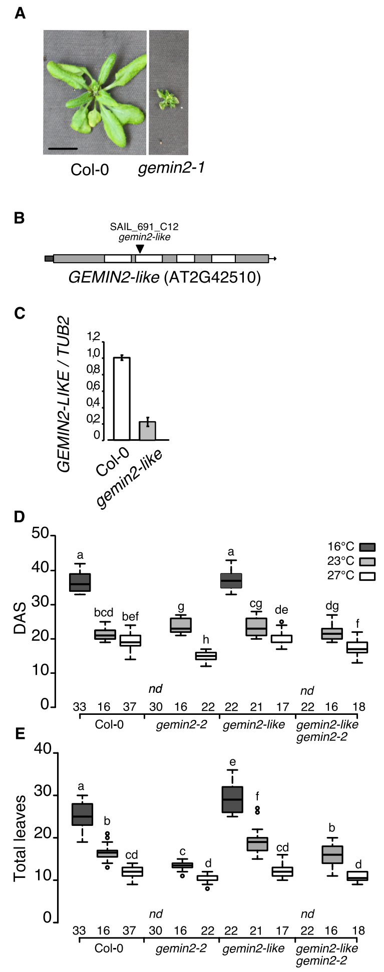


**Supplemental Figure S1. Phenotypes of *gemin2* and *gemin2 gemin2-like*.** A, Phenotype of Col-0 and *gemin2-1* grown at 16°C at 35 days-after-sowing. Scale bar: 1 cm. B, Gene structure at the *GEMIN2-like* locus. Dark-grey box indicates 5'UTR, light-grey boxes are exons, white boxes are introns. The triangle indicates the T-DNA insertion point. C, RT-qPCR analysis of *GEMIN2-* in Col-0 and *gemin2-like* mutant grown, normalized against *TUB2*. Error bars represent standard error. D,E, Flowering time of Col-0, *gemin2-2*, *gemin2like-1*, and *gemin2-2 gemin2like-1* at three different ambient temperatures. Numbers on the x-axis indicate the number of individuals. One-way ANOVA was used for statistical analysis. Different letters indicate categories that are statistically different ($p \leq 0.05$).

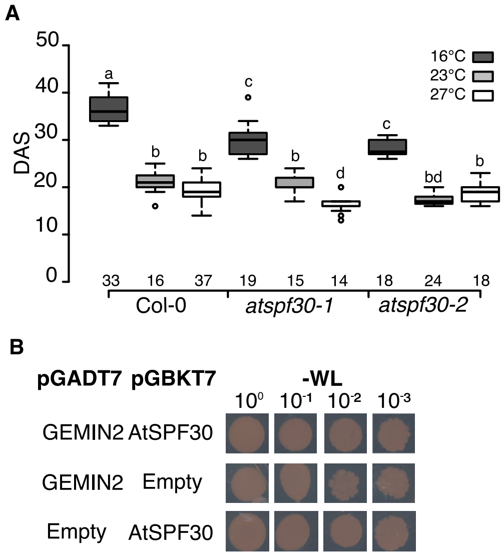


**Supplemental Figure S2.** ***AtSPF30*** **regulates flowering time.** A, Flowering time of Col-0 and two Arabidopsis *spf30* alleles at three different ambient temperatures. Numbers on the x-axis indicate the number of individuals. One-way ANOVA was used for statistical analysis. Different letters indicate categories that are statistically different ($p \leq 0.05$). B, Dilution series of yeast transformed with GEMIN2 and AtSPF30 constructs on control media (SD -WL). Photos were taken after four days after plating. pGADT7 and pGBKT7 contribute the AD and BD domains, respectively.

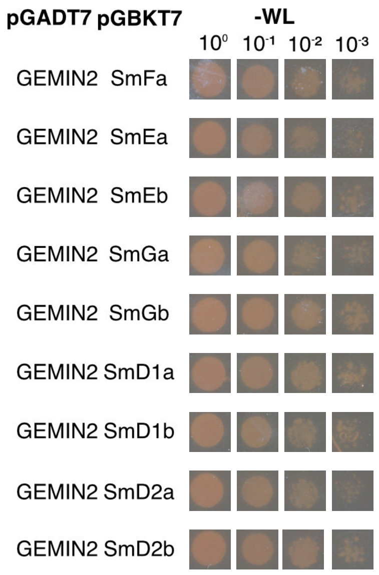


**Supplemental Figure S3. Y2H controls for GEMIN2 and Sm interaction.** Dilution series of yeast transformed with GEMIN2 and Sm constructs on control medium (SD-WL). Photos were taken after four days after plating. pGADT7 and pGBKT7 contribute the AD and BD domains, respectively.

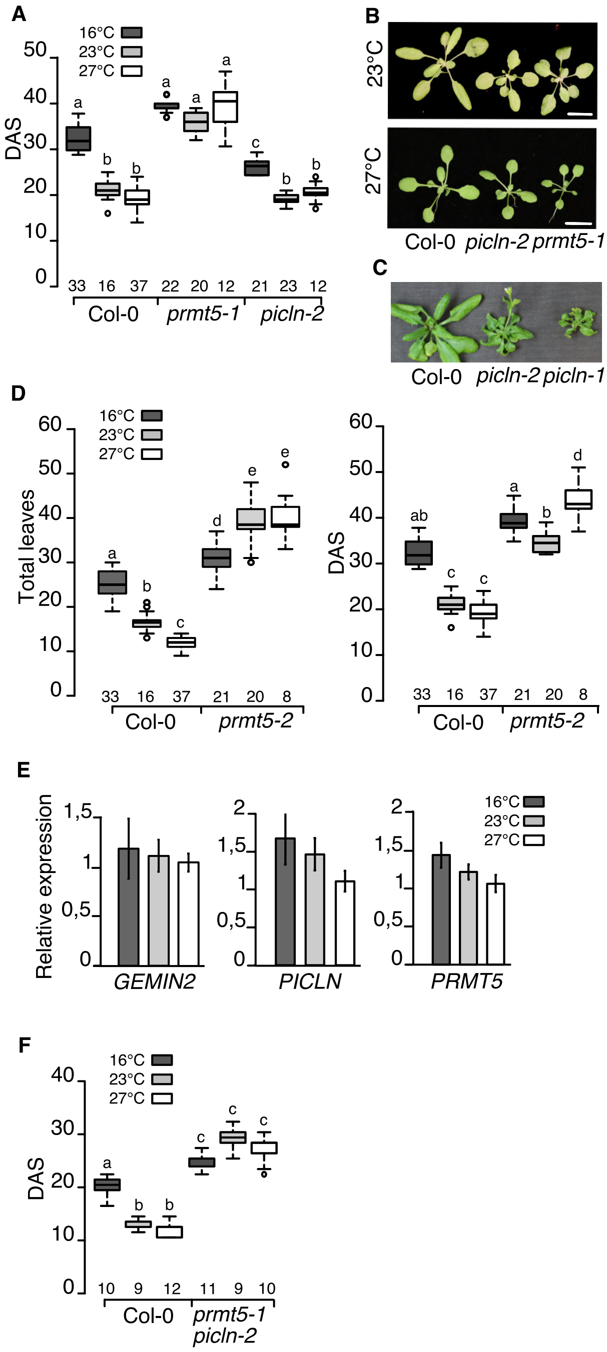


**Supplemental Figure S4. Phenotype of single and double methylosome mutants.** A, D, F, Flowering time of Col-0 and mutant genotypes at three different ambient temperatures. Numbers on the x-axis indicate the number of individuals per genotype and condition. One-way ANOVA was used for statistical analysis. Different letters indicate categories that are statistically different ($p \leq 0.05$). B, Phenotype of Col-0, *picln-2*, and *prmt5-1* grown at 23°C (top) and 27°C (bottom) for 18 and 25 days-after-sowing, respectively. Scale bar: 1 cm. C, Phenotype of Col-0, *picln-2,* and *picln-1* grown at 16°C for 35 days-after-sowing. E, RT-qPCR analysis of *GEMIN2*, *PICLN*, and *PRMT5* in Col-0 grown at three different ambient temperatures, normalized against *TUB2*. Error bars represent standard error.

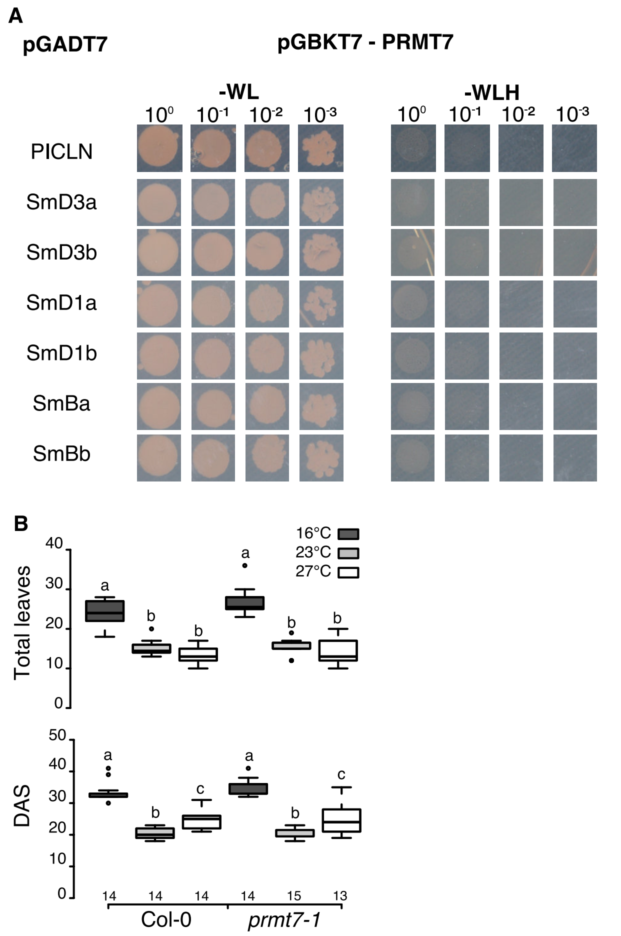


**Supplemental Figure S5. PRMT7 does not interact with PICLN or Sm proteins.** A, Y2H interaction assay between PRMT7 and PICLN or Sm proteins. Dilution series of yeast cells on control (SD-WL; left) and selective medium (SD-HWL; right). Photos were taken four days after plating. pGADT7 and pGBKT7 contribute the AD and BD domains, respectively. B, Flowering time of Col-0 and *prmt7-1* at three different temperatures.

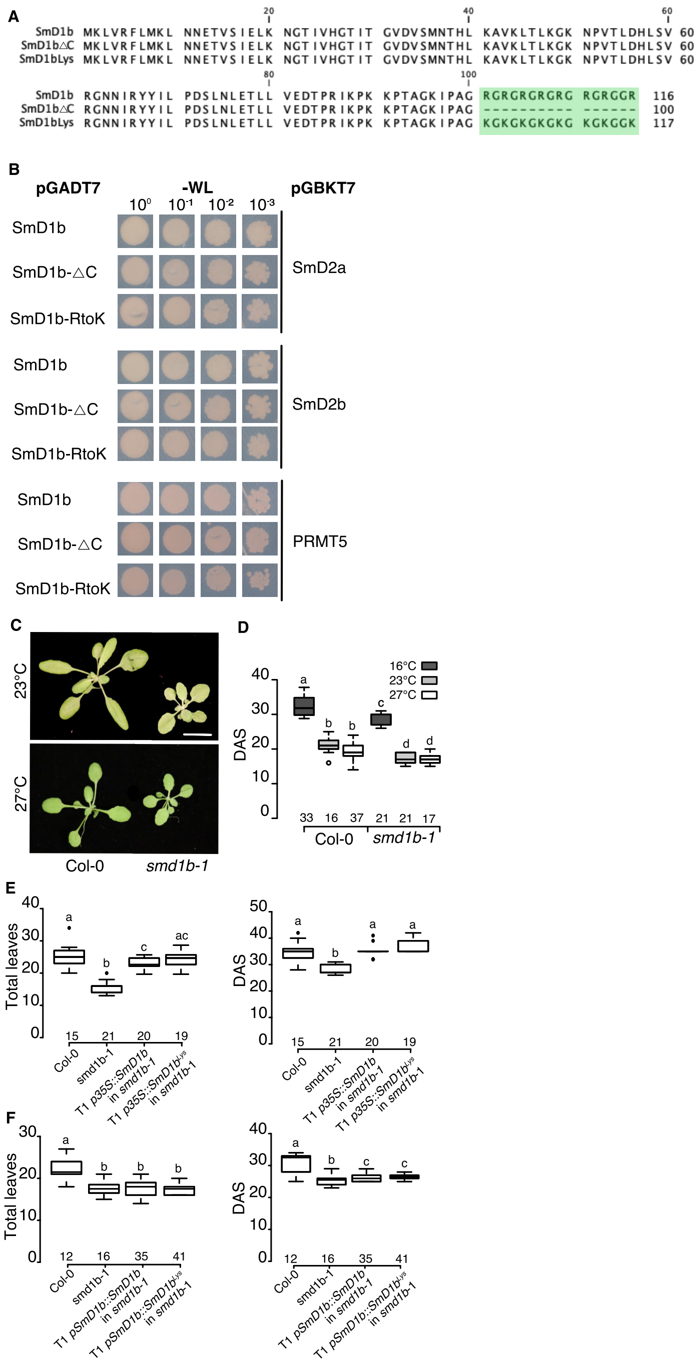


**Supplemental Figure S6. SDM is not essential for SmD1b function.** A, Alignment of SmD1b wild-type protein and mutated forms. The green box indicates the mutagenized C-term region. B, Dilution series of yeast cells grown for four days on control medium (SD-WL) for testing interaction between three different SmD1 variants and two SmD2 proteins and PRMT5. pGADT7 and pGBKT7 contribute the AD and BD domains, respectively. C, Phenotypes of Col-0 and *smd1b-1* plants grown at 23°C (top) and 27°C (bottom) for 25 and 18 DAS, respectively. D, Flowering time of Col-0 and *smd1b-1* in days-after-sowing at three different ambient temperatures. E, F, Flowering time of different T1 lines compared to wildtype and *smd1-b* mutant grown at 16°C. Numbers on the x-axis in d-f indicate the number of individuals per genotype and condition. One-way ANOVA was used for statistical analysis. Different letters indicate categories that are statistically different (p≤0.05).

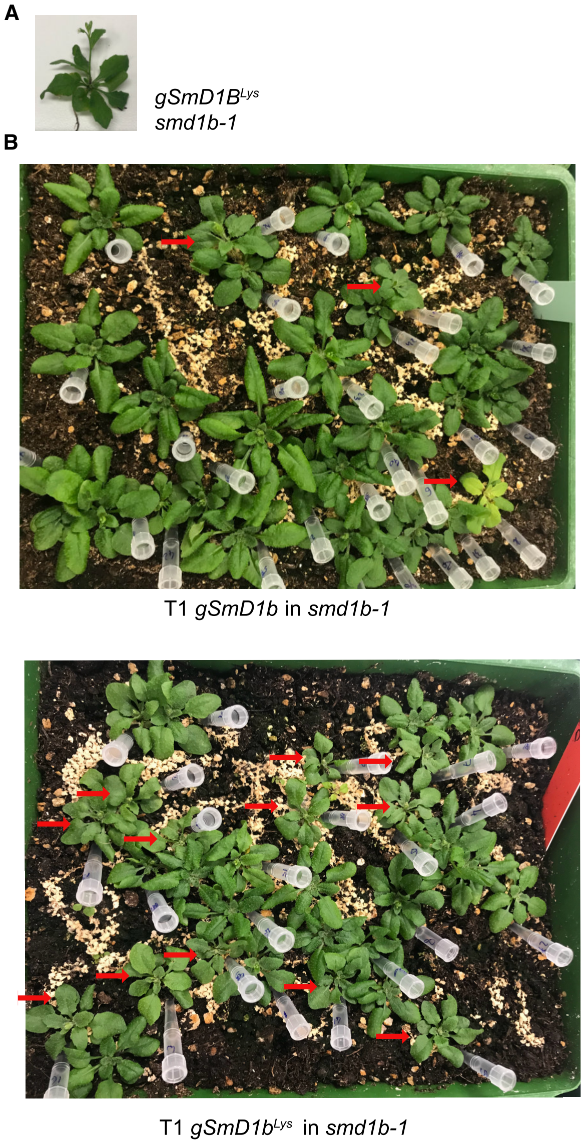


**Supplemental Figure S7. Phenotype of T1 plants expressing wild-type and mutated *SmD1b*.** A, Phenotype of *smd1b-1* transformed with gSmDb1$^{Lys}$. Plants show the *smd1b-1* mutant phenotype with irregular leaves margins and flat leaves. B, Picture of T1 plants transformed with the wild-type gSmDb1 (top) and gSmDb1$^{Lys}$ constructs. Arrows indicate plants with *smd1b-1-like* leaf phenotype.

**Supplemental Table S1. Rosette diameter (cm) taken when stem was 1 cm high.**

| **Genotype** | **16°C** | **23°C** | **27°C** |
|---|---|---|---|
| Col-0 | 8.35 ± 0.53 | 8.83 ± 0.56 | 6.59 ± 0.44 |
| *gemin2-2* | 0.88 ± 0.38 | 4.09 ± 0.43 | 6.29 ± 0.72 |
| *picln-2* | 3.38 ± 0.53 | 7.00 ± 0.71 | 5.88 ± 0.35 |
| *prmt5-1* | 7.60 ± 0.30 | 10.17 ± 0.61 | 8.73 ± 0.61 |
| *prmt5-2* | 7.10 ± 0.58 | 9.85 ± 0.46 | 9.01 ± 0.31 |
| *picln-2 prmt5-1* | 7.00 ± 0.33 | 8.89 ± 0.55 | 7.15 ± 0.82 |
| *smd1b-1* | 4.53 ± 0.56 | 6.32 ± 0.88 | 5.32 ± 0.43 |

**Supplemental Table S2. List of GEMIN2 interactors identified in Y2H screening**

| Locus Identifier | Gene Model Description | Gene Model Type | Primary Gene Symbol |
|---|---|---|---|
| AT3G60300 | RWD domain-containing protein;(source:Araport11) | protein_coding | |
| AT3G47070 | thylakoid soluble phosphoprotein;(source:Araport11) | protein_coding | |
| AT1G29880 | glycyl-tRNA synthetase / glycine-tRNA ligase;(source:Araport11) | protein_coding | |
| AT3G06150 | cytochrome P450 family protein;(source:Araport11) | protein_coding | |
| AT2G23120 | Late embryogenesis abundant protein, group 6;(source:Araport11) | protein_coding | |
| AT3G27160 | GHS1 encodes plastid ribosomal protein S21The mRNA is cell-to-cell mobile. Required for photosynthesis and C/N balance. | protein_coding | (RPS21) |
| AT1G62380 | Encodes a protein similar to 1-aminocyclopropane-1-carboxylic oxidase (ACC oxidase). Expression of the AtACO2 transcripts is affected by ethylene. | protein_coding | ACC OXIDASE 2 (ACO2) |
| AT4G34610 | Acts as a negative regulator of xylem vessel cells as a protein complex together with OVATE FAMILY PROTEIN1 (OFP1), OFP4, and MYB75. | protein_coding | BEL1-LIKE HOMEODOMAIN 6 (BLH6) |
| AT5G14740 | Encodes a beta carbonic anhydrase likely to be localized in the cytoplasm. Expression of its mRNA is seen in etiolated seedlings and points to a possible nonphotosynthetic role for this isoform. | protein_coding | CARBONIC ANHYDRASE 2 (CA2) |
| AT4G34100 | Encodes a putative E3 ubiquitin ligase that is involved in cuticular wax biosynthesis and regulates 3-hydroxy-3-methylglutaryl-CoA reductase (HMGR) activity. HMGR catalyzes the major rate-limiting step of the mevalonic acid (MVA) pathway from which sterols and other isoprenoids are synthesized. Lines carrying a recessive mutation in this locus have reduced chain-length distribution, weakly glaucous stem surface, and has reduced fertility in early flowers, non-spreading floret, downward cupped leaves, leaf waxes nearly pure C24 and C26 acid. | protein_coding | ECERIFERUM 9 (CER9) |
| AT4G23250 | cysteine-rich receptor-like protein kinase 17;(source:Araport11) | protein_coding | EMBRYO DEFECTIVE 1290 (emb1290) |
| AT5G45680 | Peptidyl-Prolyl Isomerase located in chloroplast thylakoid lumen The mRNA is cell-to-cell mobile. | protein_coding | FK506-BINDING PROTEIN 13 (FKBP13) |
| AT5G62270 | Ribosomal protein L20;(source:Araport11). Required for proper mitochondrial cristae formation. Expressed throughout plant. Mutants are defective in late stages of megagametogenesis. Pollen tube defective. Gametophytic lethality is probably due to mitochondrial disfunction. | protein_coding | GAMETE CELL DEFECTIVE 1 (GCD1) |
| AT4G08350 | global transcription factor group A2;(source:Araport11) | protein_coding | GLOBAL TRANSCRIPTION FACTOR GROUP A2 (GTA2) |
| AT2G05520 | Encodes a glycine-rich protein that is expressed mainly in stems and leaves. AtGRP3 functions in root size determination during development and in Al stress. mRNA levels are upregulated in response to ABA, salicylic acid and ethylene but downregulated in response to desiccation. The mRNA is cell-to-cell mobile. | protein_coding | GLYCINE-RICH PROTEIN 3 (GRP-3) |
| AT4G13550 | Heat stress inducible plastid monogalactosyldiacylglycerol lipase. | protein_coding | HEAT INDUCIBLE LIPASE1 (HIL1) |
| AT1G79920 | Heat shock protein 70 (Hsp 70) family protein;(source:Araport11) | protein_coding | HEAT SHOCK PROTEIN 70-15 (Hsp70-15) |
| AT5G51570 | SPFH/Band 7/PHB domain-containing membrane-associated protein family;(source:Araport11) | protein_coding | HYPERSENSITIVE INDUCED REACTION 3 (HIR3) |
| AT1G30380 | Encodes subunit K of photosystem I reaction center. The mRNA is cell-to-cell mobile. | protein_coding | PHOTOSYSTEM I SUBUNIT K (PSAK) |
| AT2G30570 | Encodes PsbW, a protein similar to photosystem II reaction center subunit W. Loss of PsbW destabilizes the supramolecular organization of PSII. | protein_coding | PHOTOSYSTEM II REACTION CENTER W (PSBW) |
| AT1G06680 | Encodes a 23 kD extrinsic protein that is part of photosystem II and participates in the regulation of oxygen evolution. Phosphorylation of this protein is dependent on calcium. In plsp1-1 mutant plastids, the nonmature form of the protein localizes in the stroma. The mRNA is cell-to-cell mobile. | protein_coding | PHOTOSYSTEM II SUBUNIT P-1 (PSBP-1) |
| AT2G43460 | Ribosomal L38e protein family;(source:Araport11) | protein_coding | RIBOSOMAL PROTEIN EL38Z (EL38Z) |

| | | | |
|---|---|---|---|
| AT3G27830 | 50S ribosomal protein L12-A The mRNA is cell-to-cell mobile. | protein_coding | RIBOSOMAL PROTEIN L12-A (RPL12-A) |
| AT5G38420 | Encodes a member of the Rubisco small subunit (RBCS) multigene family: RBCS1A (At1g67090), RBCS1B (At5g38430), RBCS2B (At5g38420), and RBCS3B (At5g38410). Activated by OXS2 under the treatment of salt. | protein_coding | RUBISCO SMALL SUBUNIT 2B (RBCS2B) |
| AT5G38410 | Encodes a member of the Rubisco small subunit (RBCS) multigene family: RBCS1A (At1g67090), RBCS1B (At5g38430), RBCS2B (At5g38420), and RBCS3B (At5g38410). Functions to yield sufficient Rubisco content for leaf photosynthetic capacity. | protein_coding | RUBISCO SMALL SUBUNIT 3B (RBCS3B) |
| AT2G47110 | polyubiquitin gene The mRNA is cell-to-cell mobile. | protein_coding | UBIQUITIN 6 (UBQ6) |

**Supplemental Table S3. List of oligos used in this study.**

| **Genotyping** | | |
|---|---|---|
| SALK_142993 (*gemin2-1*) | GTCTGTCTGTTTTCCTGCTCG | CTCATCCTCCTCTGTGTGGAG |
| SAIL_567_D05 (*gemin2-2*) | TCGAGATTACCATTGCCACTC | GTGGAATTAGCAGATATGCGG |
| SAIL_691_C12 (*gemin2like_1*) | ATTTGGAATTCGCTTAGCCTC | GCTGGTGACTGAGTACGAAGG |
| SALK_052016 (*spf30-1*) | CGGAGTTAGTGCTGAAGCAAC | TTCCAGTCACACCCACTTTTC |
| SALK_081292 (*spf30-2*) | AAAACGGCAGAGATTTTACCG | CAGCAAAGCTCAAAATCGATC |
| GABI_675D10 (*picln-1*) | TTCATCTCCATTGGAATGACC | TACTGGATGAATCTAACGGCG |
| SALK_050231 (*picln-2*) | TGGCTAAAGGATACGCAGTTG | TGAACCATCTCTTCAGCATCC |
| SALK_065814 (*prmt5-1;skb1-1*) | TGGCCTTTTGAGATGAAAGAG | GCTCATAAAGATCTTGCGCAC |
| SALK_073624 (*prmt5-2; skb1-2*) | TGGTGGAAGGAAATGGTGTAG | CTCACAGACTCTCCCATCCAG |
| SALK_039529 (*prmt7-1*) | AGAGACTGGTGTGACCATTGG | GCTAGGAAACGAACGACACAG |
| *smd1b-1* | CTGCATCTGACTTGACCAGTG | TGTGTTGTATGCTTCGTCACC |
| **Cloning** | **Foward Sequence (5' - 3')** | **Reverse Sequence (5' - 3')** |
| *GEMIN2*_promoter 1480bp (A-mod) | aacaggtctcaacctTATGCGGATCCGGTATAATC | aacaggtctcttgttGAGGCGGATGTGCGCAAAAC |
| *GEMIN2* CDS (C-mod) | aacaggtctcaggctcaacaATGATGACAAACTCCGATTC | aacaggtctctctgaCTGTCCCATTTGGCCAAAG |
| *SPF30*_CDS (C-mod) | aacaggtctcaggctcaacaATGGTAGGAGGAGTAGAAG | aacaggtctctctgaTCATTCATCAGTGCCCTCAG |
| *SMEa*_CDS (C-mod) | aacaggtctcaggctcaacaATGGCGAGCACCAAAGTTCAAAG | aacaggtctctctgaCTTTCCCGTGTTCATCATCAGAG |
| *SMEb*_CDS (C-mod) | aacaggtctcaggctcaacaATGGCGAGCACCAAAGTTCAG | aacaggtctctctgaCTTGCCCGCGTTCATCATGAG |
| *SMFa*_CDS (C-mod) | aacaggtctcaggctcaacaATGGCGTTTGTATGTATGTGTG | aacaggtctctctgatcaTTAGTCTTGATCAGCGTCTTC |
| *SMGa*_CDS (C-mod) | aacaggtctcaggctcaacaATGAGTCGTTCAGGTCAGCCTCCG | aacaggtctctctgaCTAAGAAGATCTGCCCACTGGTTCA |
| *SMD1a*_CDS (C-mod) | aacaggtctcaggctcaacaATGAAGCTCGTCAGGTTTTTG | aacaggtctctctgaACGACCTCTGCCGCGACCAC |
| *SMD1b*_CDS (C-mod) | aacaggtctcaggctcaacaATGAAGCTCGTCAGGTTTTTG | aacaggtctctctgaACGACCACCACGGCCACGTCC |
| *SMD2a*_CDS (C-mod) | aacaggtctcaggctcaacaATGAGCAAGCCAATGGAAG | aacaggtctctctgaTCATTTTGGATTCCTGAGG |
| *SMD2b*_CDS (C-mod) | aacaggtctcaggctcaacaATGAGTAAACCAATGGAAG | aacaggtctctctgaTCACTTGGGGTTCCTGAGG |
| *PRMT5*_CDS (C-mod) | aacaggtctcaggctcaacaATGCCGCTCGGAGAGAGAG | aacaggtctctctgaCTAAAGGCCAACCCAGTACG |
| *PICLN*_CDS (C-mod) | aacaggtctcaggctcaacaATGGTGGCTGGTCTAAGAG | aacaggtctctctgaCTAGTGATCTTTGGTCTCAC |

| | | |
|---|---|---|
| *PRMT7* CDS (C-mod) | aacaggtctcaggctcaacaATGTCGCCTCTGTCTTCTC | aacaggtctctctgaAGAAATAGTATGAGTGACG |
| *SMD1b*ΔC_CDS (C-mod) | aacaggtctcaggctcaacaATGAAGCTCGTCAGGTTTTTG | aacaggtctctctgaCACTAACCTGCTGGAATCTTACCTGC |
| *SMD1b*$^{Lys}$ _CDS (C-mod) | aacaggtctcaggctcaacaATGAAGCTCGTCAGGTTTTTG | aacaggtctctctgaTCATTAtttACCACCtttGCC |
| *gSMD1b* genomic (C-mod) | aacaggtctcaggctcaacaATGAAGCTCGTCAGGTATGG | aacaggtctctctgaACGACCACCACGGCCACGTC |
| *SMD1b*_promoter 1578bp (A-mod) | aacaggtctcaacctTAGTTTCAGGTTAAAGTTGATTTG | aacaggtctcttgttGGTGAGAGAATGAAGAAGAAG |
| *SMD1b*_terminator (E-mod) | aacaggtctcactgcGCATTTTGCAGTAATCAAAAG | aacaggtctcttagtATTCATCACATGCTTAGCATAAG |
| **RT-qPCR** | | |
| *GEMIN2* | AGTACCTGTTACCCATGAAG | ACATCTCCACAAGCAAATCTTC |
| *GEMIN2-like* | TCGTCTGGTATTCCTTTCTTCCA | CGGAGGAGTGTCCACTGATG |
| *PRMT5* | CTCCTGAGTGTCTTGATGGAG | TCATTGTAAAGCTTCGAAGCTG |
| *PICLN* | TCTACCCAGTTGGAGACGCTG | CCACGAACATCCATCTGATCAGC |
| *TUB2* | GAGCCTTACAACGCTACTCTGTCTGTC | ACACCAGACATAGTAGCAGAAATCAAG |

**Supplemental Table S4. List of synthesized sequences (Eurofins).**

**Hs*GEMIN2*_CDS (C-mod)**

aacaggtctcaggctcaacaATGAGAAGAGCTGAACTTGCGGGGTTGAAAACGATGGCATGGGTTCCTGCTGAATCTGCAGTGGAAGAGTTAATGCCTAGACTCTTGCCAGTA
GAGCCATGTGATCTCACTGAGGGATTTGACCCTTCAGTTCCTCCTAGAACACCGCAAGAGTACCTACGTAGGGTGCAGATTGAAGCCGCTCAATGTCCTGACGTTGTT
GTCGCTCAGATTGATCCCAAGAAGCTCAAGAGAAAACAGTCTGTGAATATCTCTCTGTCTGGATGTCAACCAGCTCCAGAGGGATATAGTCCGACCTTGCAATGGCAA
CAGCAACAGGTTGCTCAGTTCTCCACTGTTAGGCAGAATGTGAACAAACATCGATCACATTGGAAGAGCCAACAACTCGATTCAAATGTGACAATGCCGAAATCCGAA
GATGAGGAAGGTTGGAAGAAGTTCTGTTTGGGCGAGAAACTGTGTGCAGATGGTGCAGTTGGTCCAGCAACGAACGAGAGTCCTGGGATAGACTATGTGCAAATAGG
ATTTCCACCCTTATTGAGCATCGTATCGAGAATGAACCAAGCCACTGTTACCTCTGTCCTCGAGTACCTTTCGAACTGGTTTGGTGAGCGTGATTTCACACCAGAACTA
GGCAGATGGCTTTACGCGCTTCTAGCTTGCCTTGAAAAGCCTCTTCTCCCTGAAGCTCACAGTCTAATCAGGCAGTTGGCTAGGCGTTGCTCAGAGGTACGGTTGCTT
GTCGATTCCAAAGACGATGAACGAGTCCCAGCCTTAAATCTTCTGATTTGCCTTGTTTCTCGGTATTTTGACCAAAGAGATTTAGCGGATGAACCGAGCtcagagagacctgtt

**Sc*BRR1P*_CDS (C-mod)**

aacaggtctcaggctcaacaATGAAGAGAGGAGAAAGTCAAGCACCAGACGCAATCTTTGGCCAATCAAGAGCATTCGCTCTTAGTGACTCTTCGGTTAATCCTGATGTGATC
GAGTACCTTAAAAGCGTTCGACAAGAAGCACTCAGGACCAATGCGATCAGCATCAAGAACCACATGAATCTCCAGAAAAGAACTCGACATAAATCCTCTATGTATGATG
ACGAAGATGAAGGTGCCCTAAAACGTCATGCTATTTCTCCATCATTGATAAGACTGCAACGTAACGTTGAGATTTGGGTCAGATGGTTTAACTCAGTGAAAGCCACTGT
TTTAACCAATGCTTATGAATTCACAGGTTATGAAGATGAAACACTAGATCTCCTGTTATTGTTCCTTAAGAATTACCTTGAGGATATGCCGTCTAAATGTACAACGGTAGA
AAAGATCATATCTGTACTCAACCAGCATAGTTTTCCTGAGAAGGCTGAAGAGAAGGAGGAAAATCTCCAGATTGATGAGGAATGGGCTAAGAACATTTTAGTCAGGCTA
GAAAAGACGAAGATTGACTCAGTTGAGGATGTAAAGAAGGTCATAACTGAAGGAGACAAACATGAGCTGGTAGGATACAATCAGTGGTTTCAATACTTGATTAACAAC
GAACCTCAACACACTACATTTCATGAGAAAATTACGTCTAAACAGCTTTGGGTGTTGATCAAATACATGTCGAATACATGGATAAAAGAGATTCACAAGAAAGGTCGTCA
CTATAGGAGACTTCAAGATTGGTTGTTCTACATCCTTGTTCATACTCCTGAAAGAGTTACAGCGGAATATACCAGCATTCTACGGGATTTGGGGAAGAAATGCTTAGAG
CTTATCCAAAAGAAACCGGTTGAGGCTCATGAGAACAAGATTACACTTCCCAAAGAAATGGCAGAATTGAACGTCGAAATTCCAGCTGCTGTGGAGAATATGACTATCA
CCGAGTTAACTGTTTCCGTGATAGCCGTGAATTATGGGCAGAA GGATCTCATAGAGtcagagagacctgtt

***SMD1b*$^{Lys}$ CDS (C-mod)**

aacaggtctcaggctcaacaATGAAGCTCGTCAGGTTTTTGATGAAATTGAACAACGAAACAGTTTCAATCGAGCTTAAGAACGGAACCATTGTTCATGGAACCATTACTGGTG
TAGATGTTAGCATGAACACTCATTTAAAAGCTGTGAAACTCACTCTGAAAGGGAAGAATCCAGTTACATTAGACCACTTAAGTGTCAGGGGAAACAACATTCGCTATTA
TATTCTTCCGGACAGTTTGAATCTGGAGACTTTGCTGGTTGAAGACACTCCAAGAATCAAACCCAAAAAGCCAACTGCAGGTAAGATTCCAGCAGGTAAAGGAAAAG
GTAAAGGAAAAGGTAAAGGAAAAGGCAAAGGTGGTAAATAATGAtcagagagacctgtt

**g*SMD1b*$^{Lys}$_genomic (C-mod)**

aacaggtctcaggctcaacaATGAAGCTCGTCAGGTATGGTTTCTACTCGGATTCTCTTTATTTCTTCGGTTTAATTGGAATTTGCTTCGAAATTTTGAGATTTCTCTTTGTTTTCA
TGCTCTTCGATGTTTGATTTCGTATCTGGATTTTTTCGCAGGTTTTTGATGAAATTGAACAACGAAACAGTTTCAATCGAGCTTAAGAACGGAACCATTGTTCATGGAAC
CATTACTGGTACGCTTCTCTTAAAAACCCAGCTCTATTCGCTGTTTAGGGTTTTGTGTAAAATCTACGATGCTTACCTGTAATCGATGGTTACTCGCGTATTCACAAATTC
TGATGGTGTAGTTTGAGTTTAGCTAGTGTCTTTCCCCTAATAATGCTTTGTTGGTATTAACCCTAGTGACACTGGTCAAGTCAGATGCAGATGAGTACACTAGATTTTACT
TGAATCGAGGTTTATGAGATAGTTTAGTTCTTCTTCAAACTTTGATCACCCATGAGGTAGTTTACTTTCTCCTTTGATGGTTGTAACCAATGGAGCTCTCGAATTTACTAA
TTCTGATGTGTAATTCTGTTAATAGACCTAGAAATGCATTGTGGGTGTTGAATCTCTCTTGTTTGCAATAAGTGAATGATAATGGTGGTCTAGTCAGATGGAGATGATACA

TTTTATTTTGCTTTATAGTTGGGTGTATGAGAATGCTTAGTTCTTCTGGGAGTGAGTTTAGCACATGATTATATATGAGATAGTTTACTTTAGTTTCCCTCTTTATCATTGTG
GACCTCTCTTGTTAACAAATCCTGAGGCTTAATTCTGTTATTAGTGATAGTATCTGAAGAAGTTAGAGTTTAGCTGAGTCTTAATTATTTTTTTACCTCGAAATGCTTTGTT
GGTAACCCTCTTGTTAGCAAGAACTGAGTGATACTTGTCGAGTCAGATGCAGATGATTAGATTTAGCACATGAAAGTCACTCTTATCTCTTTCCTTGTAAAGAAAATCTC
TGATTTTTCACAGAACAGTTACCGTGTGGCTTCTTGTTTCAACTTCTCTTTTAGTCTGATTGATCTAATACAAGTGGTCTCTGCaGTGAGACCTGTTGCAAAATGCGCtG
TTAATGTTTGGTTTGGTTTCCTTTTTCTGCCATCATCTTGTTTAGAATGCAATTATCAATCAACTCACTGACTGGTACTATCTACTTGTGATGACTTAATGCAGGTGTAGAT
GTTAGCATGAACACTCATTTAAAAGCTGTGAAACTCACTCTGAAAGGGAAGAATCCAGTTACATTAGACCACTTAAGTGTCAGGGGAAACAACATTCGCTATTATATTCT
TCCGGACAGTTTGAATCTGGAGACTTTGCTGGTTGAAGACACTCCAAGAATCAAACCCAAAAAGCCAACTGCAGGTAACAAAACATTTCTTTTTTCTGAACTATGCCTT
TGTTTTTGTTTTTGTTTTTGCTTAGACTCATCTTTCTATTGTTCTCTTTGGTATCAGGTAAGATTCCAGCAGGTAAAGGAAAAGGTAAAGGAAAAGGTAAAGGAAAAGGC
AAAGGTGGTAAtcagagagacctgtt

**Supplemental Table S5. List of plasmids. GreenGate entry and destination vectors (Lampropoulos *et al.* 2013) were used in this study to generate the final constructs.**

| Plasmid ID | Promoter (A-mod) | N-tag (B-mod) | Insert (C-mod) | C-tag (D-mod) | Terminator (E-mod) | Selection markers (Bacteria/Plants) | Backbone |
|---|---|---|---|---|---|---|---|
| *pAD-GEMIN2* | | | GEMIN2 CDS | | | Ampicilin | pVZ22 (pGADT7 modified for GreenGate) |
| *pBD-GEMIN2* | | | GEMIN2 CDS | | | Kanamycin | pVZ23 (pGBKT7 modified for GreenGate) |
| *pBD-SPF30* | | | SPF30 CDS | | | Ampicilin | pVZ23 (pGBKT7 modified for GreenGate) |
| *pAD-SMEa* | | | SMEa CDS | | | Ampicilin | pVZ22 (pGADT7 modified for GreenGate) |
| *pBD-SMEa* | | | SMEa CDS | | | Kanamycin | pVZ23 (pGBKT7 modified for GreenGate) |
| *pAD-SMFa* | | | SMFa CDS | | | Ampicilin | pVZ22 (pGADT7 modified for GreenGate) |
| *pBD-SMFa* | | | SMFa CDS | | | Kanamycin | pVZ23 (pGBKT7 modified for GreenGate) |
| *pAD-SMGa* | | | SMGa CDS | | | Ampicilin | pVZ22 (pGADT7 modified for GreenGate) |
| *pBD-SMGa* | | | SMGa CDS | | | Kanamycin | pVZ23 (pGBKT7 modified for GreenGate) |
| *pAD-SMGb* | | | SMGb CDS | | | Ampicilin | pVZ22 (pGADT7 modified for GreenGate) |
| *pBD-SMGb* | | | SMGb CDS | | | Kanamycin | pVZ23 (pGBKT7 modified for GreenGate) |
| *pAD-SMD1a* | | | SMD1a CDS | | | Ampicilin | pVZ22 (pGADT7 modified for GreenGate) |
| *pBD-SMD1a* | | | SMD1a CDS | | | Kanamycin | pVZ23 (pGBKT7 modified for GreenGate) |
| *pAD-SMD1b* | | | SMD1b CDS | | | Ampicilin | pVZ22 (pGADT7 modified for GreenGate) |
| *pBD-SMD1b* | | | SMD1b CDS | | | Kanamycin | pVZ23 (pGBKT7 modified for GreenGate) |
| *pAD-SMD2a* | | | SMD2a CDS | | | Ampicilin | pVZ22 (pGADT7 modified for GreenGate) |
| *pBD-SMD2a* | | | SMD2a CDS | | | Kanamycin | pVZ23 (pGBKT7 modified for GreenGate) |
| *pAD-SMD2b* | | | SMD2b CDS | | | Ampicilin | pVZ22 (pGADT7 modified for GreenGate) |
| *pBD-SMD2b* | | | SMD2b CDS | | | Kanamycin | pVZ23 (pGBKT7 modified for GreenGate) |

| | | | | | | | |
|---|---|---|---|---|---|---|---|
| *pAD-PRMT5* | | | PRMT5 CDS | | | Ampicilin | pVZ22 (pGADT7 modified for GreenGate) |
| *pBD-PRMT5* | | | PRMT5 CDS | | | Kanamycin | pVZ23 (pGBKT7 modified for GreenGate) |
| *pAD-PICLN* | | | PICLN CDS | | | Ampicilin | pVZ22 (pGADT7 modified for GreenGate) |
| *pBD-PICLN* | | | PICLN CDS | | | Kanamycin | pVZ23 (pGBKT7 modified for GreenGate) |
| *pAD-PRMT7* | | | PRMT7 CDS | | | Ampicilin | pVZ22 (pGADT7 modified for GreenGate) |
| *pBD-PRMT7* | | | PRMT7 CDS | | | Kanamycin | pVZ23 (pGBKT7 modified for GreenGate) |
| *pAD-SMD1bΔC* | | | *SMD1bΔC* CDS | | | Ampicilin | pVZ22 (pGADT7 modified for GreenGate) |
| *pBD-SMD1bΔC* | | | *SMD1bΔC* CDS | | | Kanamycin | pVZ23 (pGBKT7 modified for GreenGate) |
| *pAD-SMD1b$^{Lys}$* | | | *SMD1b$^{Lys}$* CDS | | | Ampicilin | pVZ22 (pGADT7 modified for GreenGate) |
| *pBD-SMD1b$^{Lys}$* | | | *SMD1b$^{Lys}$* CDS | | | Kanamycin | pVZ23 (pGBKT7 modified for GreenGate) |
| *pGEMIN2::GEMIN2CDS* | pGEMIN2 | B-dummy | GEMIN2 CDS | D-dummy | tRBCS | Spec/Basta | pGGZ003 |
| *pGEMIN2::HsGEMIN2* | pGEMIN2 | B-dummy | HsGEMIN2 CDS | D-dummy | tRBCS | Spec/Basta | pGGZ003 |
| *pGEMIN2::ScBRR1P* | pGEMIN2 | B-dummy | ScBRR1P CDS | D-dummy | tRBCS | Spec/Basta | pGGZ003 |
| *p35S::SMD1bCDS* | p35S | B-dummy | SMD1b CDS | D-dummy | tRBCS | Spec/Basta | pGGZ003 |
| *p35S::SMD1b$^{Lys}$CDS* | p35S | B-dummy | *SMD1b$^{Lys}$* CDS | D-dummy | tRBCS | Spec/Basta | pGGZ003 |
| *pSMD1b::SMD1bCDS::tSMD1b* | pSMD1b | B-dummy | SMD1b CDS | D-dummy | tSMD1b | Spec/Basta | pGGZ003 |
| *pSMD1b::SMD1b$^{Lys}$CDS::tSMD1b* | pSMD1b | B-dummy | *SMD1b$^{Lys}$* CDS | D-dummy | tSMD1b | Spec/Basta | pGGZ003 |
| *pSMD1b::gSMD1b::tSMD1b* | pSMD1b | B-dummy | gSMD1b | D-dummy | tSMD1b | Spec/Basta | pGGZ003 |
| *pSMD1b::gSMD1b$^{Lys}$::tSMD1b* | pSMD1b | B-dummy | *gSMD1b$^{Lys}$* | D-dummy | tSMD1b | Spec/Basta | pGGZ003 |
